# A Novel Humanized Mouse Model for HIV and Tuberculosis Co-infection Studies

**DOI:** 10.1101/2024.03.05.583545

**Authors:** José Alejandro Bohórquez, Sitaramaraju Adduri, Danish Ansari, Sahana John, Jon Florence, Omoyeni Adejare, Gaurav Singh, Nagarjun Konduru, Chinnaswamy Jagannath, Guohua Yi

**Author notes:** Authors contributed equally to this work (José Alejandro Bohórquez, Sitaramaraju Adduri).

## Abstract

Tuberculosis (TB), caused by *Mycobacterium tuberculosis* (*Mtb*), continues to be a major public health problem worldwide. The human immunodeficiency virus (HIV) is another equally important life-threatening pathogen. Further, co-infections with HIV and *Mtb* have severe effects in the host, with people infected with HIV being fifteen to twenty-one times more likely to develop active TB. The use of an appropriate animal model for HIV/*Mtb* co-infection that can recapitulate the diversity of the immune response in humans would be a useful tool for conducting basic and translational research in HIV/*Mtb* infections. The present study was focused on developing a humanized mouse model for investigations on HIV-*Mtb* co-infection. Using NSG-SGM3 mice that can engraft human stem cells, our studies showed that they were able to engraft human CD34+ stem cells which then differentiate into a full-lineage of human immune cell subsets. After co-infection with HIV and *Mtb*, these mice showed decrease in CD4+ T cell counts overtime and elevated HIV load in the sera, similar to the infection pattern of humans. Additionally, *Mtb* caused infections in both lungs and spleen, and induced the development of granulomatous lesions in the lungs, detected by CT scan and histopathology. Distinct metabolomic profiles were also observed in the tissues from different mouse groups after co-infections. Our results suggest that the humanized NSG-SGM3 mice are able to recapitulate the effects of HIV and *Mtb* infections and co-infection in the human host at pathological, immunological and metabolism levels, providing a dependable small animal model for studying HIV/*Mtb* co-infection.

## INTRODUCTION

Tuberculosis (TB) remains one of the biggest public health problems worldwide, being the second cause of death in mankind in 2022, behind COVID-19 ^1^. Over seven million people were newly diagnosed with TB in the past year and around 1.3 million people were killed by this deadly disease. There is a consensus that a quarter of the world population are infected with *Mycobacterium tuberculosis* (*Mtb)*, the causative agent for TB^1^. The majority of *Mtb*-infected individuals remain latently infected without clinical signs (LTBI). However, around 10% of the infected patients will develop active TB, especially in conjunction with immunodeficiency caused by malnutrition, immunosuppressive therapy using steroids, or infection with immunosuppressive pathogens^2^. Among these infection with human immunodeficiency virus (HIV) plays a pivotal role, given that immunosuppression is the hallmark of HIV pathogenesis^3^. HIV is the etiological agent for acquired immunodeficiency syndrome (AIDS), another equally important public health concern responsible for the death of over 40 million people as of 2023^4^. The synergy between HIV and *Mtb* in co-infection has been extensively examined, and compelling evidence showed that HIV exacerbates TB severity, and is the leading cause of death for people infected with *Mtb*^4–6^. This is likely because HIV-induced immunosuppression leads to a disruption of CD4 T cells, the main driver of Th-1 immunity in LTBI patients, resulting in active TB^7^.

Non-human primates (NHP) are routinely used as large animal models for HIV/*Mtb* research not only because the monkeys and humans have remarkably similar genomes, physiology, and immune systems, but also because the monkeys can be infected by both *Mtb* and Simian immunodeficiency virus (SIV)^8^. The latter is also a retrovirus and belongs to the same Lentivirus genus as HIV and causes HIV-like infection in NHPs. After co-infection, NHPs also display AIDS-like features as in humans, such as massive reduction of CD4+ T cells and a high viral load in the sera if without anti-retroviral treatment, as well as chronic immune activation in animals during extended observation ^7, 9^. Furthermore, the co-infected monkeys also recapitulate key aspects of human TB infection stages, including latent infection, chronic progressive infection, and acute TB, depending on the route and dose of infection^10–12^. Importantly, *Mtb* latently infected macaques co-infected with SIV results in reproducible LTBI reactivation^13^, providing a reliable model for HIV/*Mtb* research. However, NHPs require specialized infrastructure for experimentation and are cost-restrictive, and are not readily available in the majority of animal facilities^14, 15^,

The use of other small animal models, such as rodents poses different challenges. Although inbred and genetic knockout mice are easily available, and readily infective using *Mtb*, most strains of mice are not a natural host for HIV, which require human CD4^+^ T cells to establish infection. Whereas the use of mouse models for *Mtb* research has also been criticized due to their inability to form granulomas which are a hallmark of *Mtb* infection in humans^16^, certain mouse strains and infection protocols show the formation of granulomas. ^17^. Fortunately, humanized mouse models, the immunodeficient mice that have been reconstituted with a human immune system, appears to be a promising small animal model for HIV and *Mtb* reseach^14, 15, 18, 19^. They have been extensively used for evaluating HIV gene therapy and therapeutics^20, 21^, and recently, the NSG (NOD scid gamma)-based humanized BLT mice were developed for analyzing *Mtb* and HIV/*Mtb* co-infections^15, 18, 22^. However, humanized BLT mice need surgical transplantation (under the kidney capsule) of fetal liver, bone marrow and thymus tissues, and restriction of human fetal tissues used for research and the sophisticated surgery has markedly limited the use of this model. In addition, these mice have immature B cells with poor IgG class-switching and poor reconstitution of myeloid lineage of antigen-presenting cells (APCs)^23, 24^, posing a challenge for HIV/*Mtb* research because myeloid cells, especially macrophages, are important targets for both HIV and *Mtb*.

We demonstrate here that these deficiencies can be ameliorated in the newly developed NSG-SGM3 mice, which transgenically express three human cytokine/chemokine genes IL-3, GM-CSF, and KITLG. The expression of these genes improves the differentiation and maturation of the myeloid cells^25–29^. The present study is aimed at establishing a reliable new-generation, humanized mouse model for the HIV/*Mtb* co-infection research. We show that humanized NSG-SGM3 mice can differentiate CD34+ stem cells into a full-lineage of immune cell subsets, including both lymphoid and myeloid lineages. Importantly, we show that HIV/*Mtb* infections are reproducible in these mice with a spectrum of immunological, pathological, and metabolic changes when compared to uninfected mice.

## MATERIALS AND METHODS

### Bacterial and viral strains

*Mtb* H37Rv was obtained from BEI Resources (USA) and propagated in the biosafety level 3 (BSL-3) facilities at the University of Texas Health Science Center at Tyler (UTHSCT). It was cultured in 7H9 broth with 10% OADC supplement following standard *Mtb* culture procedures^30^. After 7 days of growth, the bacteria were collected and subjected to sonication three times, at an amplitude of 38%, for 10 seconds/each, with a 5-second interval, followed by low-speed centrifugation (1,100 RPM). Bacteria were diluted to an optical density (OD) value of ≈ 1 in sterile NaCl 0.9% and aliquots were made and frozen at -80 °C to be used as inoculum. Two weeks later, one aliquot was thawed, and the bacterial content was evaluated by plating ten-fold serial dilutions in 7H10 agar, supplemented with OADC. After 3 weeks of incubation, the colony forming units (CFU) per mL were calculated.

HIV-1 BaL strain was obtained from NIH AIDS Reagent Program, also prepared in the BSL-3 facilities at UTHSCT, following standard procedures ^31^. Briefly, frozen human PBMCs (STEMCELL Technologies, Vancouver, Canada) were thawed and seeded in a 75 cm^2^ flask at a concentration of 5 × 10^6^ cells/mL in RPMI 1640 media (Corning Inc., Corning, NY) supplemented with 10% fetal bovine serum (FBS), 1% penicillin/streptomycin, 1 µg/ml of PHA and 2 µg/ml polybrene (MilliporeSigma, Burlington, MA). After 3 days of stimulation, 4 × 10^7^ cells were centrifuged and infected with HIV-1 BaL using an MOI (multiplicity of infection) of 0.1 (4 × 10^6^ TCID_50_) in two adsorption cycles. Following the second adsorption cycle, the cells were seeded in two 75 cm^2^ flasks with 30 ml of media supplemented with FBS, antibiotics, and human IL-2 (20 Units/ml). Cell culture supernatant was collected every three days, with fresh media being added, until day 21 of culture and stored at -80 °C. A small aliquot from each collection will be used to titrate the virus using quantitative RT-PCR.

### Animal experiment design

All animal procedures were approved by the UTHSCT Institutional Animal Care and Use Committee (IACUC) (Protocol #707). NOD.Cg-Prkdc^scid^ Il2rg^tm1Wjl^ Tg(CMV-IL3,CSF2,KITLG)1Eav/MloySzJ (NSG-SGM3) mice were purchased from The Jackson laboratory (Bar Harbor, ME) and bred in the Vivarium facilities at UTHSCT. Pups were weaned at 21 days after birth and, 1-3 weeks after that, they were irradiated at a dose of 100 cgy/mouse, followed by intravenous injection with 2 × 10^5^ CD34^+^ stem cells/mouse at 12 h post-irradiation. Humanization was monitored starting at 12 weeks after stem cell transplantation and again at 14 and 16 weeks. For this purpose, blood was drawn from the submandibular vein (100-150 µl, based on animal weight) and PBMCs were collected through density gradient centrifugation using Ficoll Paque (Cytiva, Marlborough, MA). After erythrocyte lysis, the PBMC from each animal were stained for human (hu) and mouse (mo) hematopoietic cell surface marker (CD45^+^), as well as lymphocytic and myeloid markers. Animals that showed a positive huCD45^+^/moCD45^+^ ratio, accompanied by differentiation of various immune cell populations, were selected for experimental infection.

Mice were randomly divided into four experimental groups: Uninfected (n=5), HIV-infected (n=8), *Mtb*-infected (n=8) and HIV/*Mtb* co-infected (n=7). *Mtb* infection was performed using aerosolized *Mtb* H37Rv through a Madison chamber, as previously described^32^, using an infection dose of 100 CFU/mouse. Three additional mice were included in the Madison chamber at the time of infection and were euthanized 24 hours after infection. The lungs were collected, macerated and plated on 7H10 agar, to confirm the initial bacterial implantation^33^.

One day after *Mtb* infection, the mice for the HIV alone and HIV/*Mtb* co-infection groups were subjected to intraperitoneal (IP) inoculation with 10^5^ TCID_50_ of HIV_BaL_. Blood samples from all experimental groups were collected on the day of infection and at 15-, 28- and 35-days post infection (dpi). Serum samples from all the animals were separated and stored at -80 °C until further use. PBMCs were isolated and stained for flow cytometry analysis. At 35 dpi, the animals were terminally anesthetized, using a Ketamine/Xylazine mixture, in order to perform computed tomography (CT) scan and pulmonary function (PF) tests. Afterwards, the animals were euthanized and whole blood samples were collected through cardiac punction. During necropsy, lung and spleen samples were collected and macerated through a 70 μM cell strainer (Thermo Fisher scientific) in a final volume of 2 ml of PBS. Serial ten-fold dilutions of the organ macerates were plated in 7H10 agar, supplemented with OADC, to assess the bacterial load. The remaining volume of lung and spleen macerates were stored at -80 °C for further analysis.

For each experimental group, lung sample from one animal was selected for histopathological analysis and, therefore, not subjected to maceration and bacterial culture. Lungs were filled with 10% formalin, before being removed from the animal, and stored in the same media after the necropsy^34, 35^. Sample processing and Hematoxylin-Eosin (HE) staining was carried out at the histopathology core of UT southwestern.

### CT scan and PF testing

Mice were intraperitoneally injected with ketamine/xylazine (100 mg/kg Ketamine, 20 mg/kg Xylazine). Once the correct anesthetic plane was achieved, the mice were intubated with a sterile, 20-gauge intravenous cannula through the vocal cords into the trachea. Following intubation, anesthesia was maintained using isoflurane.

Elastance (Ers), compliance (Crs), and total lung resistance (Rrs) was assessed for each mouse through the snapshot perturbation method, as previously described^36^. Measurements were performed in triplicates for each animal, using the FlexiVent system (SCIREQ, Tempe, AZ), with a tidal volume of 30 mL/kg at a frequency of 150 breaths/min against 2–3 cm H2O positive end-expiratory pressure.

After PF testing, the mice were subjected to CT scans for the measurements of lung volume, using the Explore Locus Micro-CT Scanner (General Electric, GE Healthcare, Wauwatosa, WI). CT scans were performed during full inspiration and at a resolution of 93 μm. Lung volumes were calculated from lung renditions collected at full inspiration. Microview software 2.2 (http://microview.sourceforge.net) was used to analyze lung volumes and render three-dimensional images.

### RNA extraction and RT-qPCR

Serum samples from all experimental groups were extracted using the NucleoSpin RNA isolation kit (Macherey-Nagel, Allentown, PA). Following viral RNA extraction, samples were evaluated using RT-qPCR to determine the viral RNA load in each animal^37^. Control standards (obtained from NIH AIDS Reagent Program) with known quantities of HIV-1 genome copies were used as amplification controls, as well as to stablish a standard curve that was used to determine the viral RNA load, based on the cycle threshold (Ct) value.

### Flow cytometry analysis

Flow cytometry was performed using the PBMCs from all experimental animals at the specified sampling timepoints. In all cases, the PBMCs isolated from each animal were divided into two wells of a 96-well U-shaped bottom plate (Corning Inc., Corning, NY), used for staining with two separate flow cytometry panels. Cells were washed and inoculated with Fc block (Biolegend, San Diego, CA) at 4 °C for 20 minutes, followed by another wash. Afterwards, cells were incubated with fluorescence-conjugated monoclonal antibodies. For the first flow cytometry panel, cells were incubated with antibodies against the following human surface markers: Alexa Fluor™ 405-CD45, FITC-CD3, APC-CD4, PE-CD8, PerCP-CD56, Alexa Fluor™ 510-CD19 (Biolegend, San Diego, CA). For the second flow cytometry panel, the antibodies against human cell surface markers were as follows: Alexa Fluor™ 405-CD45, Alexa Fluor™ 510-CD86, APC-CD11b, PE-CD11c, PerCP-HLA-DR, Alexa Fluor™ 700-CD14 (Biolegend, San Diego, CA). Additionally, for the second panel, the cells were also incubated with an FITC-labelled antibody against moCD45. After staining, the cells were washed and fixed for 1 hour, followed by another wash. Flow cytometry was performed using the Attune NxT flow cytometer (Invitrogen, Waltham, MA), including the corresponding isotype controls for each antibody. Analysis was carried out with the FlowJo software v10.6.1 (BD life sciences), using the isotype controls as guidelines for gating.

### Immunofluorescence staining

Paraffin-embedded lung sections were used for immunofluorescent staining against human immune cell subsets^38^. Samples were deparaffined by submerging the slides in Xylene (Fisher bioreagents), followed by sequentially lower concentrations of ethanol. Afterwards, antigen retrieval and blocking of non-specific binding were performed, using 10mM sodium citrate buffer and PBS with 0.4% triton and 5% FBS, respectively. Primary antibody incubation was carried out overnight at 4 °C with human-CD68 monoclonal antibody (cat. No. 14-0688-82, Invitrogen) and CD19 Rabbit polyclonal antibody (cat. No. 27949-1-AP, Proteintech, Rosemont, USA), diluted in PBS + 0.4% triton + 1% FBS at the recommended dilutions. The following day, samples were incubated for 2 hours at room temperature with goat anti-mouse IgG1-Alexa Fluor™ 568 (cat. No. A21124, Invitrogen) and goat anti-rabbit IgG-Alexa Fluor™ 488 (cat. No. A11008, Invitrogen), at the recommended dilutions. The slides were mounted using DAPI-supplemented mounting medium (Abcam, Cambridge, UK) and images were captured with a LionheartLX automated microscope (Biotek, Winoovski, VT). Images were processed with the GEN5 software version 3.09 (Biotek) and the ImageJ software (NIH).

### Multiplex assay for cytokine profiling

The cytokine profile in lung and spleen tissue macerate, as well as serum samples at 35 dpi, from all experimental groups were evaluated in duplicates using the Bio-Plex Pro™ Human Cytokine panel (Bio-Rad, Hercules, CA), according to the manufacturer’s instructions. Briefly, 50 µL of filtered tissue homogenate, or 1:4 diluted serum, were dispensed in a 96-well plate containing magnetic beads conjugated with antibodies for the detection of 27 different cytokines. Following incubation with detection antibodies and streptavidin-PE, the samples were analyzed in the Bio-Plex MAGPIX multiplex reader (Bio-Rad Laboratories Inc., CA). A regression curve, based on the values obtained from a set of standard dilutions, was used to convert the fluorescence values reported by the machine into cytokine concentrations (expressed as pg/mL).

The 27 cytokines and chemokines reported by the Bio-Plex Pro™ Human Cytokine panel were: Basic FGF, Eotaxin, G-CSF, GM-CSF, IFN-γ, IL-1β, IL-1Ra, IL-2, IL-4, IL-5, IL-6, IL-7, IL-8, IL-9, IL-10, IL-12, IL-13, IL-15, IL-17, IP-10, MCP-1, MIP-1α, MIP-1β, PDGF-BB, RANTES, TNF-α and VEGF.

### Mouse blood sample handling for metabolomic analysis

Whole blood sample was collected from mice in all the experimental groups at the end of the study and plasma was separated through centrifugation. The samples were processed for collection of the metabolite pellet as follows: 50 μl of plasma were mixed with 950 μl of 80% ice-cold methanol, followed by centrifugation at >20.000 G for 15 minutes in a refrigerated centrifuge. Afterwards, the supernatant was transferred to a new tube and vacuum dried, using no heat. The metabolite pellet was analyzed at the metabolomic core facility at the Children’s Medical Center Research Institute at University of Texas Southwestern Medical Center (Dallas, TX, USA) using liquid chromatography–mass spectrometry (LC-MS), as previously described^39^.

### Metabolome data analysis

Statistical analysis of metabolome profiles was performed in R environment (R version 4.1.0). Raw abundance values of metabolites were used as input for statistical analysis. The raw data was log2 transformed and normalized across the samples using ‘limma’ package^40^ by cyclically applying fast linear loess normalization with a 0.3 span of loess smoothing window and 10 iterations wherein each sample was normalized to pseudo-reference sample which was computed by averaging all samples. Principal components analysis was performed using ‘PCAtools’ package. Orthogonal partial least squares discriminant analysis (OPLS-DA) was performed and variable importance on projection (VIP) score were computed using ‘ropls’ package. VIP score of >1 is considered for feature selection. Hierarchical clustering was performed on normalized data after univariate scaling. Hierarchical clustering was performed using correlation to calculate clustering distance with averaging method for clustering. Differentially abundant metabolites (DAMs) were identified using student t test. The correlation between metabolite abundances and *Mtb* or HIV loads were analyzed using Pearson correlation method. For all hypothesis testing analyses, statistical significance was set 5% (p value = 0.05) to reject null hypothesis.

### Statistical analysis

Statistical differences between groups were assessed using the Prism software version 8.3.0. for Windows (GraphPad Software, San Diego, California USA, www.graphpad.com). Unpaired, non-parametric, t-tests were employed for different comparisons between groups.

## RESULTS

### Human CD34+ HSCs-engrafted NSG-SGM3 mice can differentiate a full array of human immune cell phenotypes

After 16 weeks of humanization, PBMCs from the hCD34^+^ HSCs-transplanted mice were evaluated by flow cytometry for human lymphoid and myeloid cell surface markers. The NSG-SGM3 mice allow stem cells to develop into human lymphoid lineages, such as T cells (CD3^+^, between 10-90%, including both CD4+ and CD8+ T cells) and B cells (CD19^+^, between 7-60%) (**Fig. 1**). Additionally, differentiation of human myeloid subsets (CD14^+^) was also observed, ranging between 1 and 25%. Within the myeloid lineage, we also detected CD11b+ macrophages (**Fig. 1**, Gating strategy is shown in **Supplementary Fig. 1**).

**Figure 1.**
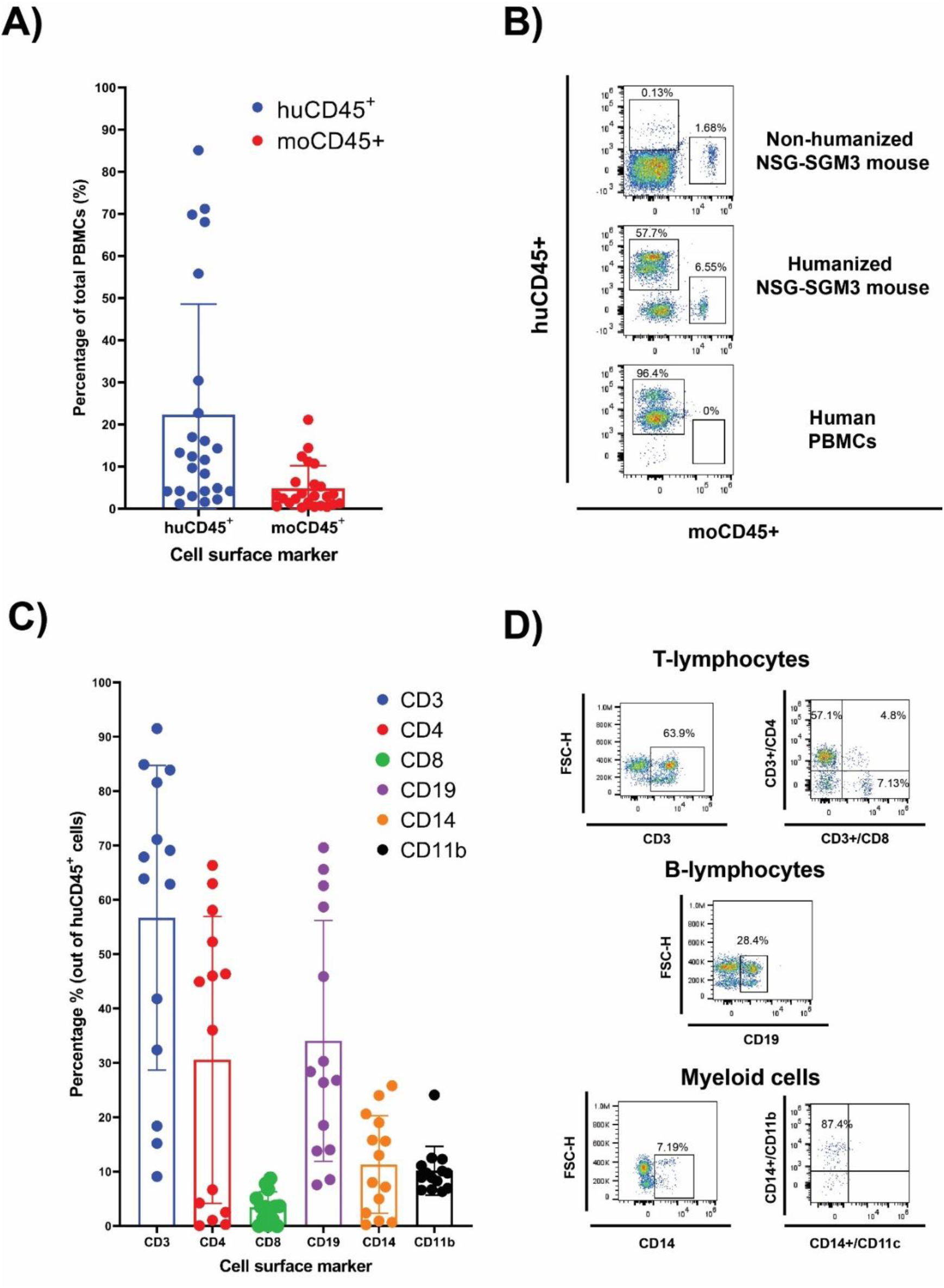
Human CD34^+^ hematopointci stem cells (HSC) engraftment and differentiation of human immune cells in the NSG-SGM3 mice. **(A and B)**: The differential expression of humanCD45 (huCD45) and mouseCD45 (moCD45) expressing cells in mice after 14 weeks of humanization. Percentages of human and mouse CD45**^+^** cells are shown as histogram in A (n=27), and the representative flow cytometry dot plot of the comparative expression of human cell surface markers between the humanized NSG-SGM3 mice and human PBMCs are shown in B. **(C)** Percentages of human immune cell populations (n=27). **(D)** representative flow cytometry dot plot of T lymphocytes, B cells and myeloid cells.

### Humanized NSG-SGM3 mice are susceptible to both HIV-1 and *Mtb* infections

After HIV/*Mtb* infections, HIV viral RNA was detected in serum samples from the infected mice starting at 15 dpi, with most animals in the HIV single-infection group being positive at this time, while only two out of the seven mice in the HIV/*Mtb* co-infection group showed viral RNA (**Fig. 2a**). The viral RNA load detected in the positive animals at 15 dpi was between 2×10^5^ and 2.2×10^6^ copies/ml. However, all the HIV-infected animals were positive in subsequent samplings at 28 and 35 dpi. The HIV RNA load was between 3.7×10^4^ and 6.8×10^5^ copies/ml for animals with single HIV infection and between 4.1×10^4^ and 7.7×10^5^ copies/ml for the HIV/*Mtb* co-infected mice. No significant differences were detected in the viral RNA load between the two HIV-infected groups at these timepoints.

**Figure 2.**
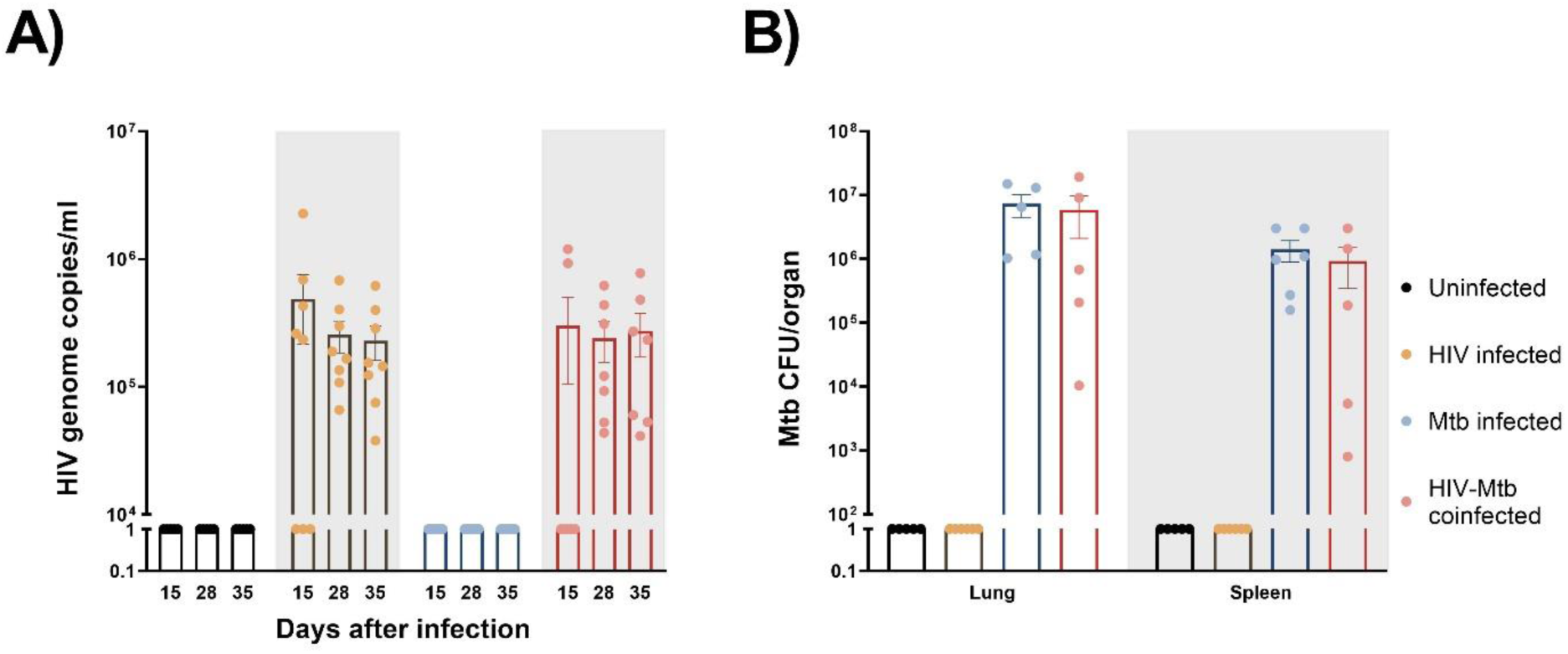
Establishment of HIV-1 and *Mycobacterium tuberculosis (Mtb)* infections in humanized mice. **(A)** HIV-1 RNA load, expressed as genome copies/mL, was assessed in serum samples from all experimental groups at three different timepoints of the study **(B)** *Mtb* bacterial load in lungs and spleens, expressed as CFU/organ, was evaluated in all experimental groups at the end of the study.

The *Mtb* bacterial load was assessed in lung and spleen samples after euthanasia in the *Mtb* single infection group and the HIV/*Mtb* coinfected mice (**Fig. 2b**). In both groups, a higher bacterial load was found in lungs than in spleens. Moreover, the mean CFU count in the lungs and spleens from *Mtb* single infection group (7.3×10^6^ and 1.4×10^6^, respectively) was higher than the animals co-infected with HIV (5.8×10^6^ for lung and 9.2×10^5^ for spleen), even though their differences are not significant (**Fig. 2b**).

### Immune phenotype changes in humanized mice after infection

We also monitored the human immune cell population changes over time after HIV/*Mtb* infections. Starting from 15 dpi, huCD45^+^/moCD45^+^ ratio was significantly decreased (p<0.05) in the two HIV-infected groups (HIV single infection and HIV/*Mtb* co-infection), and the huCD45^+^/moCD45^+^ ratio decrease was sustained until the late stage of the experiment. Conversely, the *Mtb* single infection group showed similar or even increased huCD45^+^/moCD45^+^ ratio after infection (**Fig. 3a**).

**Figure 3.**
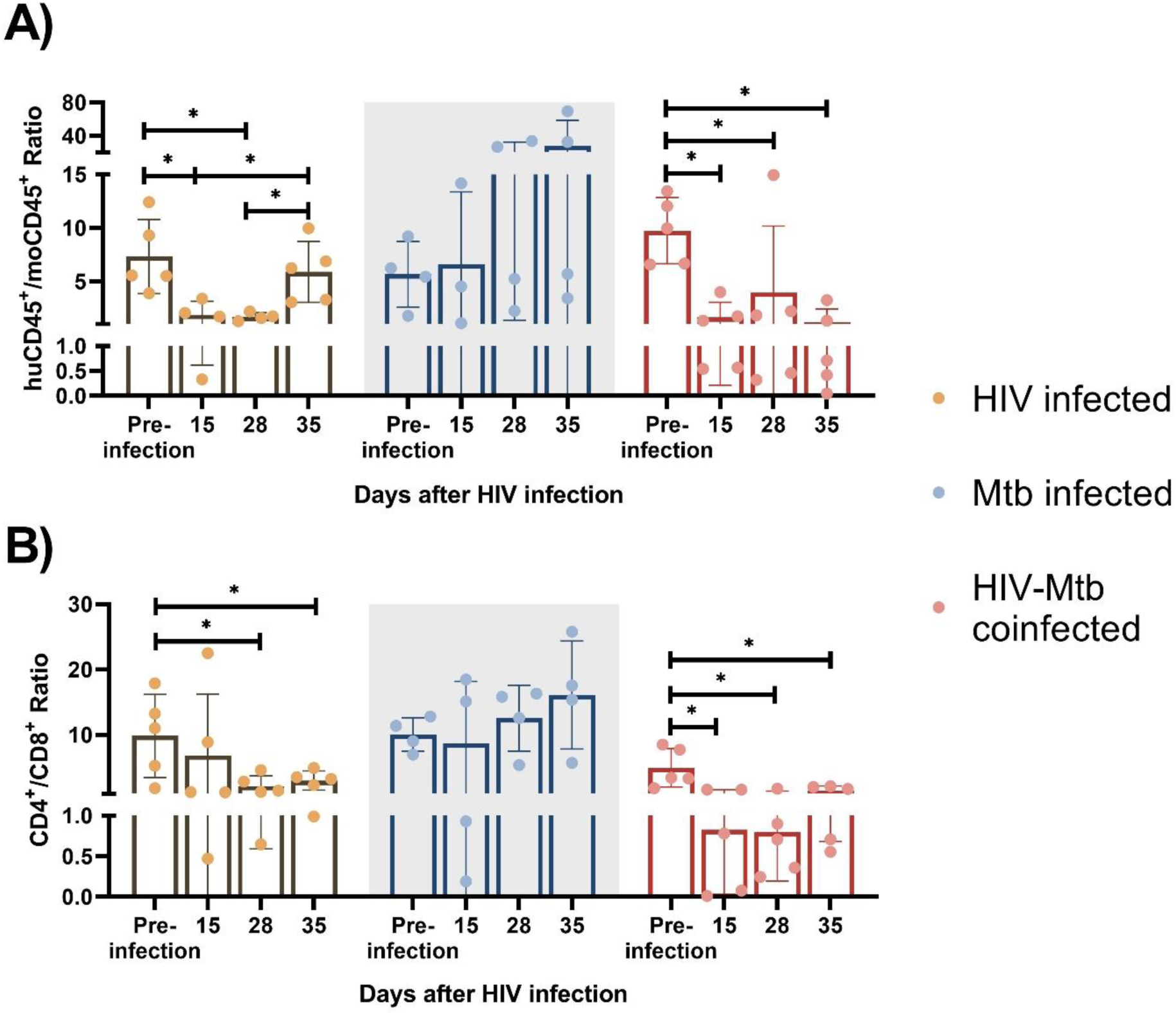
Immune cell phenotype changes after HIV-1 and *Mtb* infections. HuCD45^+^/moCD45^+^ ratio **(A)** and CD4^+^/CD8^+^ ratio **(B)** were calculated for each infected animal at different timepoints after infection. Asterisk indicates statistically significant differences (p<0.05, what test)

We observed significant CD4^+^ T cell depletion in the HIV-infected groups (HIV single infection and HIV/*Mtb* co-infection). We used CD4^+^/CD8^+^ ratio as an indicator for CD4^+^ T cell depletion, and we found a ∼10-fold CD4^+^/CD8^+^ reduction (p<0.05) in the HIV/*Mtb* co-infected mice as early as 15 dpi, and this trend remained until the end of the experiment. In the single infection group, we also found a lower mean CD4^+^/CD8^+^ ratio since 15 dpi, while the subsequent samplings at 28 and 35 dpi showed significant decreases on CD4^+^/CD8^+^ ratio values. In contrast, there was no significant difference detected over time in the Mtb alone infection group (**Fig. 3b**).

### Alterations in cytokines and chemokines production in humanized mice after infection

In serum sample, significant increases in G-CSF, MCP-1 and MIP-1α was detected in the *Mtb* single infection group, in comparison with both HIV-infected groups (**Fig. 4a**). Additionally, the serum concentration of IL-2 and IL-8 were also significantly increased in the *Mtb* single infection group, compared to the HIV/*Mtb* co-infection. The HIV/*Mtb* co-infected mice analyzed showed higher IP-10 than both the HIV and *Mtb* single infection mice (**Fig. 4a**).

**Figure 4.**
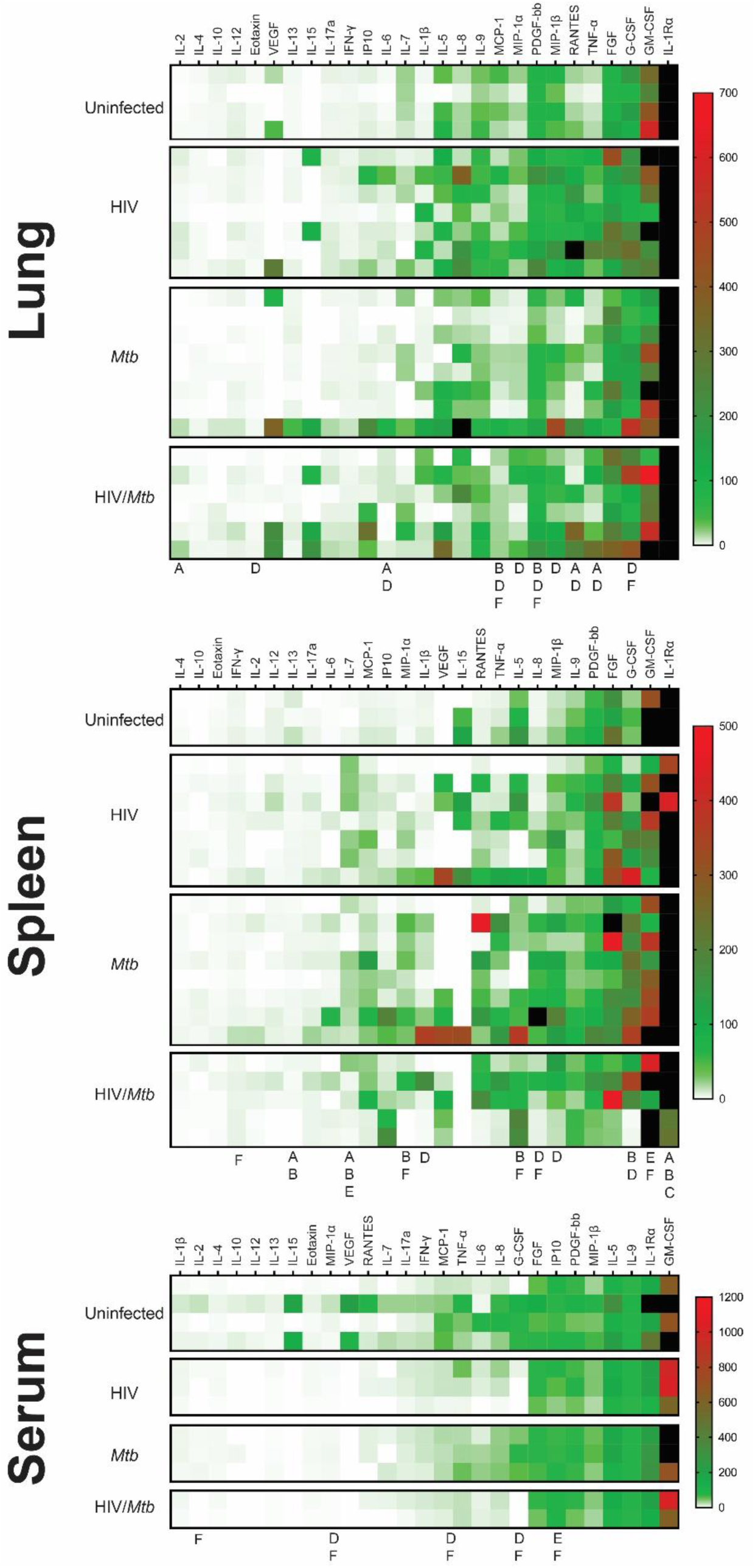
Cytokine profiles (Heatmap) in serum, lung and spleen samples. The Bio-Plex Pro™ Human Cytokine panel was used in the multiplex assay to evaluate the concentrations of 27 different human cytokines, which are expressed as pg/ml. **(A)** Cytokine profile of lung samples. **(B)** Cytokine profile of spleen samples. **(C)** Cytokine profile of serum samples. the letters under the columns show differences as follows: **A**: Difference between uninfected and HIV-infected, **B**: Difference between uninfected and *Mtb*-infected, **C**: difference between uninfected and HIV/*Mtb*-coinfected, **D**: Difference between HIV-infected and *Mtb*-infected, **E**: difference between HIV-infected and HIV/*Mtb*-coinfected, and **F**: Difference between *Mtb*-infected and HIV/*Mtb*-coinfected. (p<0.05; unpaired T test). Note: The black color on the right of heatmap shows the far high value that are out-of-range levels.

Lung macerate supernatants showed an increase in the concentration of IL-6, RANTES and TNF-α in the HIV single infection group compared to the uninfected control animals, as well as the Mtb single infection group (**Fig. 4b**). Additionally, IL-2 concentrations were also higher in the HIV-infected animals than in the uninfected mice. Moreover, HIV single infection also induced statistically higher levels of Eotaxin, MIP-1α and MIP-1β than single *Mtb* infection. Statistical analysis also revealed a decrease in MCP-1 and PDGF concentration in lung samples from *Mtb* infected mice, compared to the remaining three experimental groups (**Fig. 4b**).

In the case of spleen samples, macerates from the *Mtb* single-infection group were found to have significantly higher concentrations of IL-1β, G-CSF and MIP-1β than the HIV single-infection group (**Fig. 4c**). Similarly, the levels of IL-8 and MIP-1α were higher in the *Mtb* group than in both HIV-infected groups. In contrast, both the HIV and *Mtb* single infection groups showed lower concentrations of GM-CSF than the HIV/*Mtb* co-infected animals, while this group also had statistically higher amounts of IFN-γ than the *Mtb* group. All the infected groups showed a decrease in IL-1Rα and IL-13, compared to the uninfected control animals (**Fig. 4c**).

### *Mtb* infection induced pathological changes in the lungs of humanized mice

We stained the lung section with H&E staining, and we observed diffuse immune cell infiltration in lung sample from *Mtb*-infected mice. In some cases, immune cell infiltration was observed around a necrotic nucleus, in structures similar to TB granulomas. No such cellular aggregates were detected in either the uninfected or the HIV single infection groups (**Fig. 5a**). We stained lung sections from *Mtb*-infected humanized mice by immunofluorescent staining, and the result showed that the cell populations surrounding the necrotic area mostly corresponded with macrophages (CD68^+^), though other immune cell types, such as CD19^+^ B cells, were also found. However, no granuloma structure was observed in the lung section of the uninfected mice, even though a low proportion of cells expressing the human CD68^+^ and CD19^+^ surface markers was observed in the lung sections from uninfected mice (**Fig. 5b**).

**Figure 5.**
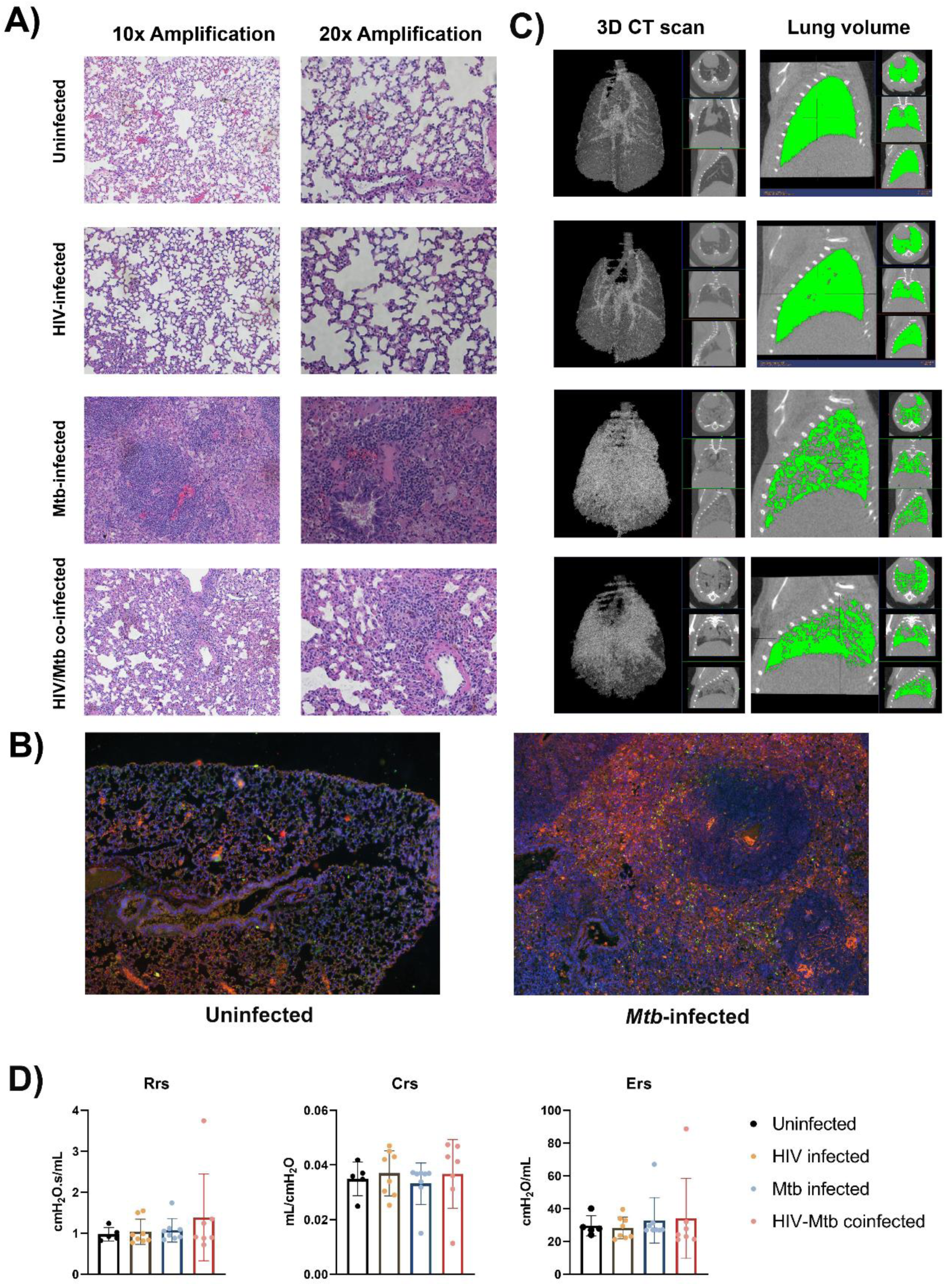
Histopathological, radiological and functional changes in the lungs of NSG-SGM3 mice after HIV/*Mtb* infection and coinfection. **(A)** Lung sections were obtained from formalin-fixed tissues of animals in all experimental groups (one animal for each group) and subjected to hematoxylin-eosin staining, two different amplifications are shown. **(B)** Immunofluorescence staining of surface markers for human macrophages (CD68-Alexafluor 568, in orange) and B-cells (CD19-Alexa 488, in green) in lung sections from uninfected and *Mtb*-infected mice. DAPI-supplemented mounting buffer (in blue) was used for nuclei staining. **(C)** Representative 3D renditions of CT scan and lung volume pictures obtained from animals in all experimental groups. **(D)** Pulmonary function test parameters: Resistance (Rrs), compliance (Crs) and elastance (Ers), were collected from animals in all experimental groups at the end of the trial (Uninfected: n=5; HIV-infected: n=8; *Mtb*-infected: n=8; HIV/*Mtb*-co-infected: n=7).

The CT scan showed an increase in high density areas in the *Mtb*-infected animals, regardless of their HIV-infection status, indicating the occurrence of inflammation and other pathological changes in the lungs (**Fig. 5c**). However, no significant differences were detected in the pulmonary function tests between the experimental groups (**Fig. 5d**).

### Different plasma metabolome landscapes in healthy mice, HIV infection, *Mtb* infection and co-infection

Plasma metabolome profiling was performed for a total of 10 samples including no infection (n=3), *Mtb* infection (n=3), HIV infection (n=2), and HIV/*Mtb* co-infection (n=2). Abundances of 175 metabolites were estimated. To enable comparison of metabolite abundances between different samples, data was normalized across the samples. To investigate differences in plasma metabolome landscape among the four categories of infection, principal components analysis (PCA) was performed. PCA is an unsupervised learning method suitable for dimensionality reduction of high dimensional metabolome data. Interestingly, the plasma metabolome profiles are stratified according to infection status in PCA (**Fig. 6a**). Mice with no infection appeared distinct from all infected mice. While the mice with infections were clustered separately from healthy mice, there was a clear distinction among HIV infection alone, *Mtb* infection alone, and HIV/*Mtb* co-infection. This suggests that the global plasma metabolome is distinctly altered based on infection status and type. Interestingly, the samples from HIV/*Mtb* co-infected mice clustered in between HIV infection alone and *Mtb* infection alone suggesting they show metabolic changes common for individual infections.

**Figure 6.**
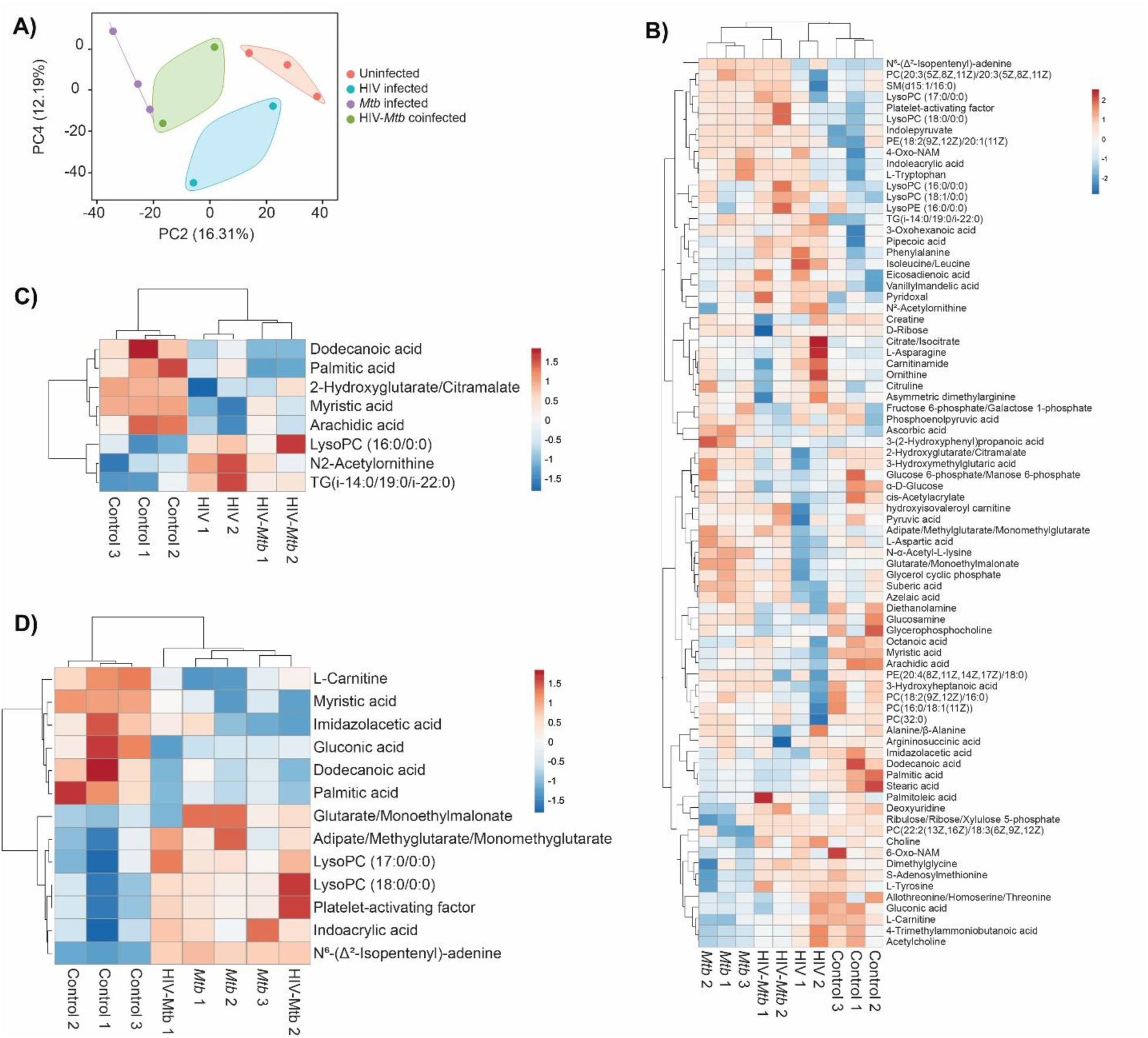
Metabolomics analysis of the plasma from healthy and HIV and/or *Mtb*-infected humanized mice. **(A)** Principal components analysis of plasma metabolome profiles of mice from no infection (n=3), *Mtb* infection (n=3), HIV infection (n=2), and dual infection (n=2) categories. Two principal components were selected to plot a two-dimensional graph to depict variation across the sample categories. Variance explained by each of the two components was given in parenthesis. **(B)** Heatmap showing abundances of 75 metabolites with a VIP score > 1 computed in OPLS-DA on plasma metabolome profiles of mice from no infection (n=3), *Mtb* infection (n=3), HIV infection (n=2), and dual infection (n=2) categories. Normalized data was scaled using univariate scaling. Hierarchical clustering was performed using correlation to calculate clustering distance with averaging method for clustering. **(C** and **D)** Heatmap showing differentially abundant metabolites in (**C**) HIV infection and (**D**) *Mtb* infection compared with healthy mice. Normalized data was scaled using univariate scaling. Hierarchical clustering was performed using correlation to calculate clustering distance with averaging method for clustering.

To identify metabolites varying across the four categories, we performed OPLS-DA followed by computation of VIP scores on all 175 metabolites. OPLS-DA is a supervised analysis which helps in identifying variables that discriminate different categories of samples based on VIP score. There were 75 metabolites with a VIP score >1 (**Supplementary Table 1**). The abundances of these metabolites across all four categories were shown with hierarchical clustering (an unsupervised algorithm) in **Fig. 6b**. As expected, in concordance with PCA, dendrogram of hierarchical clustering showed that infection and no infection categories are distinct, while co-infection stratified between the two individual infections (**Fig. 6b**).

To identify metabolites that are differentially abundant in HIV infection, we compared healthy mice (n=3) to HIV infection mice (n=4; HIV infection alone and HIV/*Mtb* co-infection). We identified 8 DAMs in HIV infection with a p value <0.05 (**Fig. 6c** and **Table 1**). Similarly, we compared healthy mice (n=3) to *Mtb* infection mice (n=5; *Mtb* infection alone and HIV/*Mtb* co-infection) to identify metabolites differentially abundant in *Mtb* infection which yielded 13 DAMs (**Fig. 6d** and **Table 2**). Interestingly, three fatty acids, namely dodecanoic acid, palmitic acid and myristic acid exhibited reduced abundance in HIV infected mice as well as *Mtb* infection mice (**Table 1** and **2**).

**Table 1:**
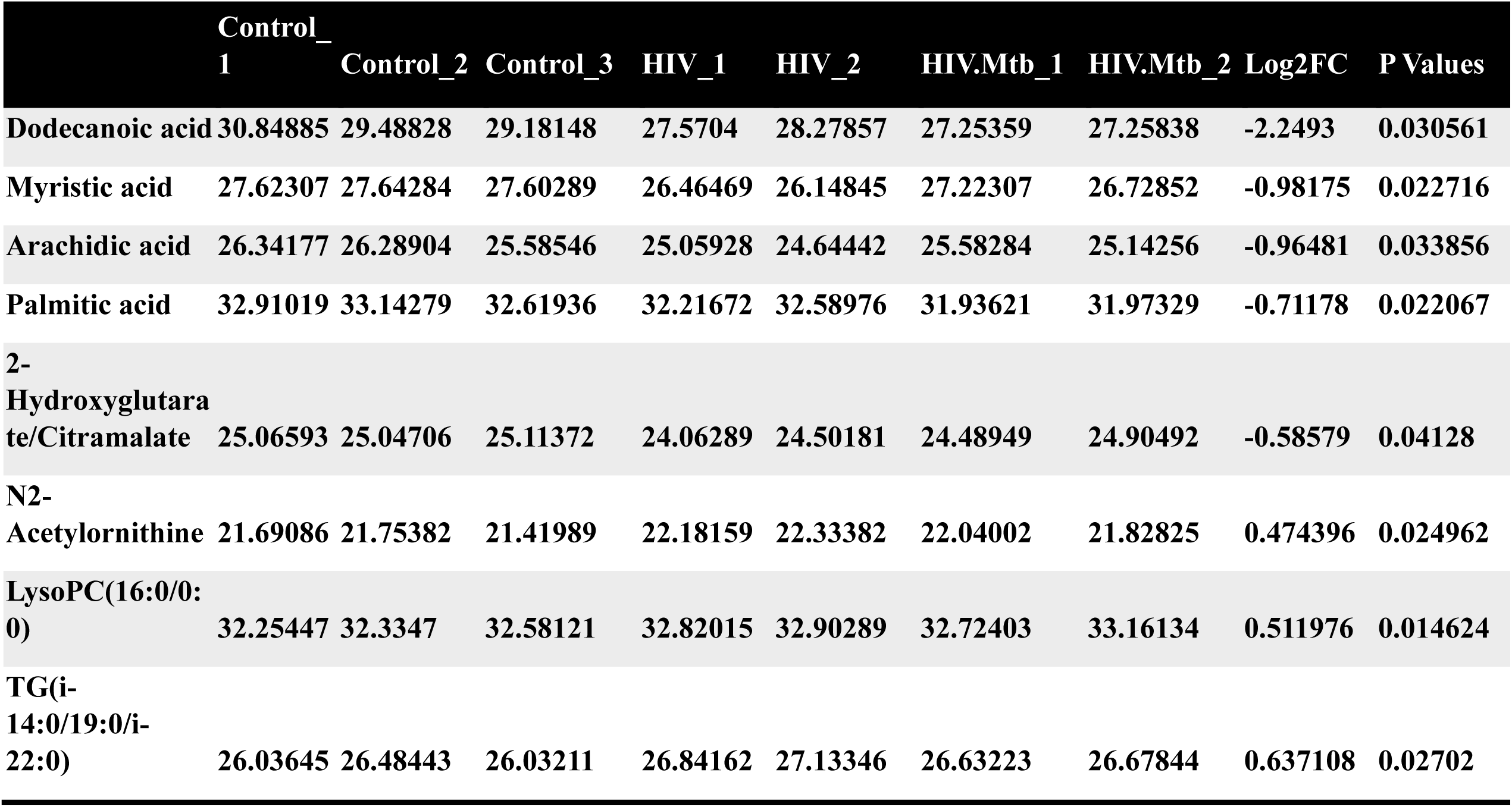
Metabolites differentially abundant in HIV infection.

|  | Control_1 | Control_2 | Control_3 | HIV_1 | HIV_2 | HIV.Mtb_1 | HIV.Mtb_2 | Log2FC | P Values |
| --- | --- | --- | --- | --- | --- | --- | --- | --- | --- |
| Dodecanoic acid | 30.84885 | 29.48828 | 29.18148 | 27.5704 | 28.27857 | 27.25359 | 27.25838 | -2.2493 | 0.030561 |
| Myristic acid | 27.62307 | 27.64284 | 27.60289 | 26.46469 | 26.14845 | 27.22307 | 26.72852 | -0.98175 | 0.022716 |
| Arachidic acid | 26.34177 | 26.28904 | 25.58546 | 25.05928 | 24.64442 | 25.58284 | 25.14256 | -0.96481 | 0.033856 |
| Palmitic acid | 32.91019 | 33.14279 | 32.61936 | 32.21672 | 32.58976 | 31.93621 | 31.97329 | -0.71178 | 0.022067 |
| 2-Hydroxyglutarate/Citramalate | 25.06593 | 25.04706 | 25.11372 | 24.06289 | 24.50181 | 24.48949 | 24.90492 | -0.58579 | 0.04128 |
| N2-Acetylornithine | 21.69086 | 21.75382 | 21.41989 | 22.18159 | 22.33382 | 22.04002 | 21.82825 | 0.474396 | 0.024962 |
| LysoPC(16:0/0:0) | 32.25447 | 32.3347 | 32.58121 | 32.82015 | 32.90289 | 32.72403 | 33.16134 | 0.511976 | 0.014624 |
| TG(i-14:0/19:0/i-22:0) | 26.03645 | 26.48443 | 26.03211 | 26.84162 | 27.13346 | 26.63223 | 26.67844 | 0.637108 | 0.02702 |

**Table 2:**
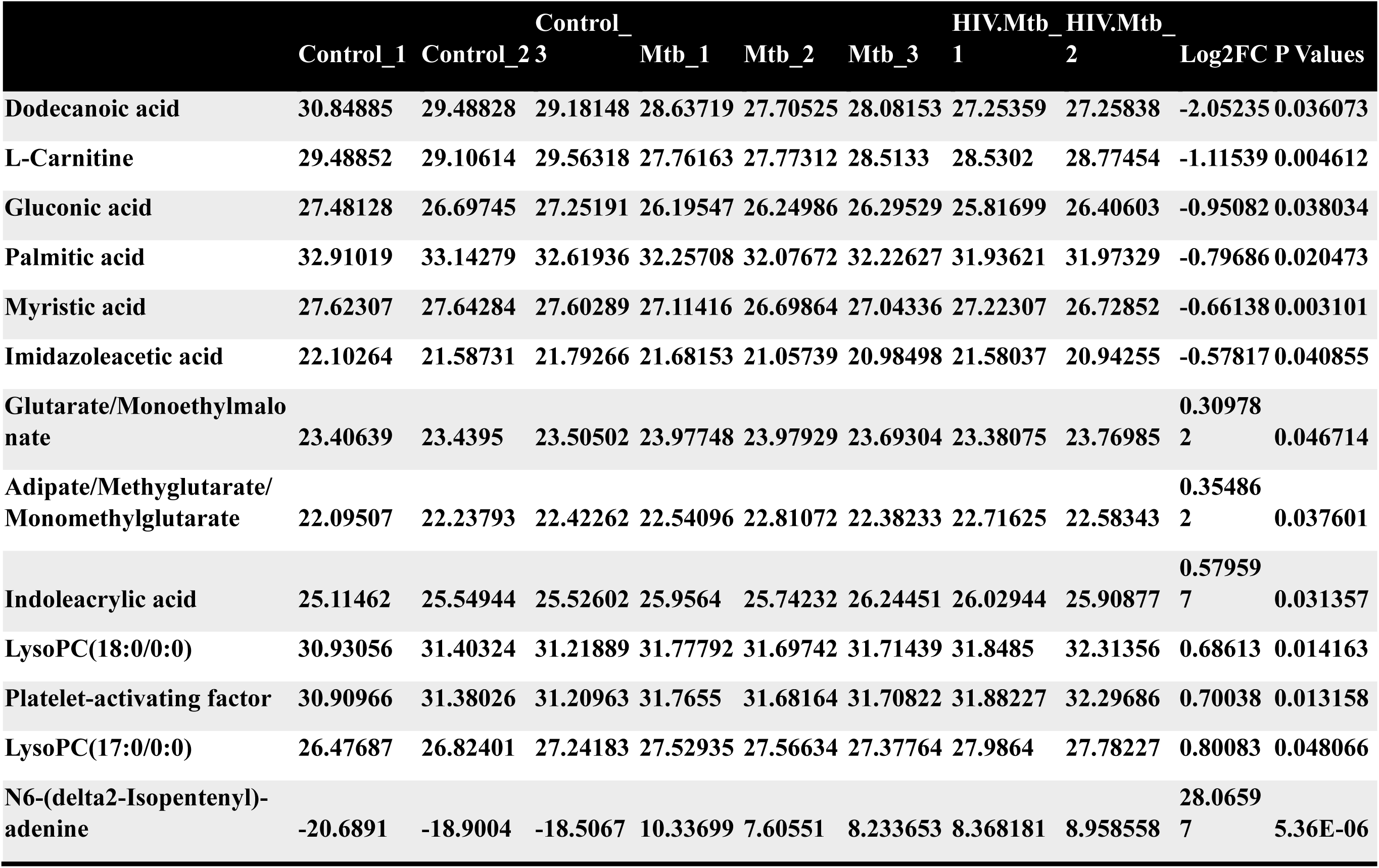
Metabolites differentially abundant in Mtb infection.

### Metabolite abundances correlated with HIV and *Mtb* loads

To identify metabolites correlating with HIV or *Mtb* load with metabolites, we used Pearson correlation analysis. HIV infection load (as detected by RNA copies/ml plasma) positively correlated with diethanolamine (r=0.99), and negatively correlated with glucose 6-phosphate/mannose 6-phosphate (r=-0.95) and imidazole acetic acid (r=-0.92) (**Fig. 7A**).

**Figure 7.**
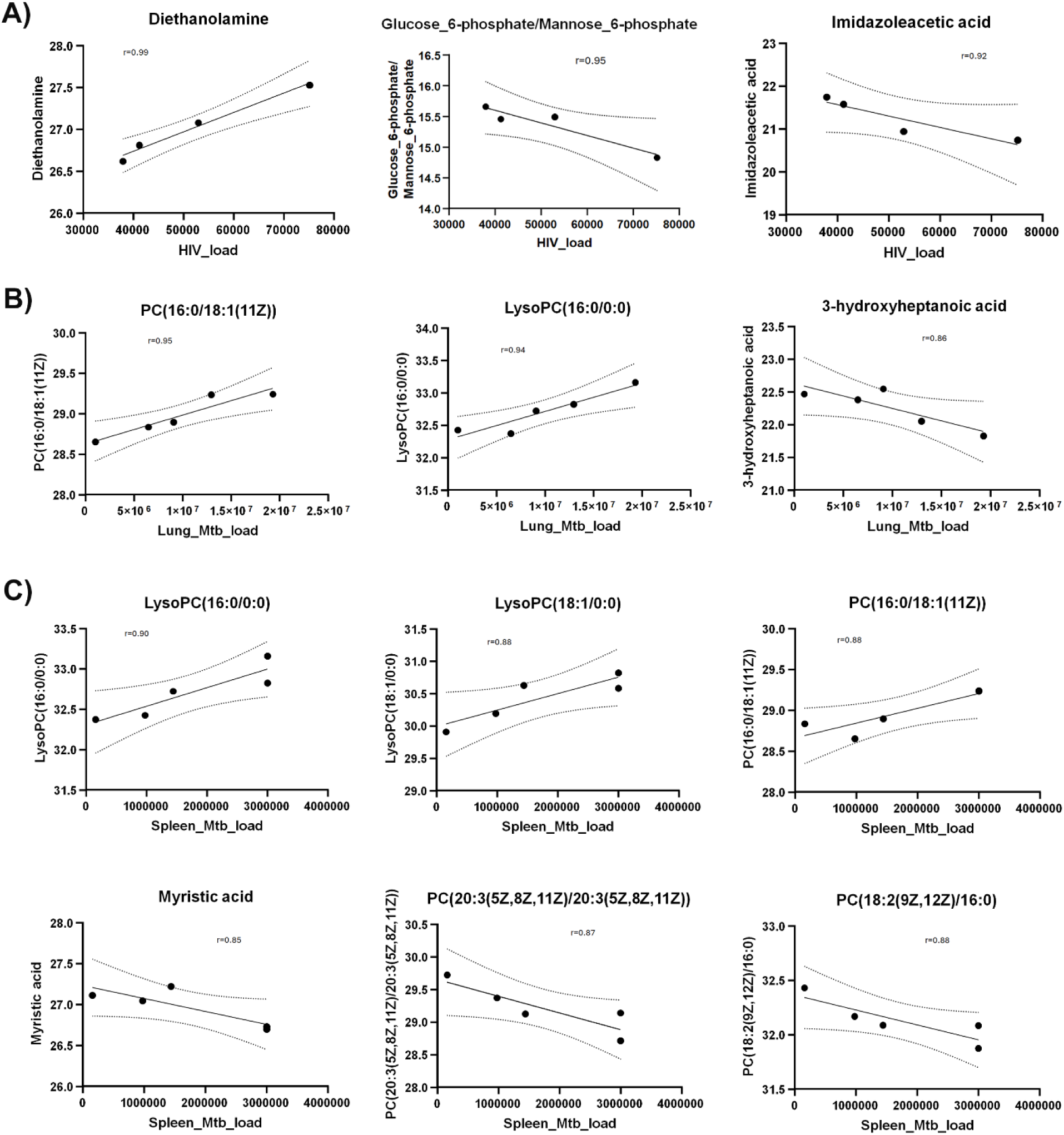
Scatter plots show Pearson correlation between metabolites and HIV/*Mtb* load in mice. **(A)** Pearson correlation between metabolites and serum HIV load (viral copies/ml). **(B)** Pearson correlation between metabolites and *Mtb* load in lungs (CFU/lung). **(C)** Pearson correlation between metabolites and *Mtb* load in spleens (CFU/spleen). Y axis shows normalized metabolites abundance values. Dotted curves show 95% confidence interval of model fit. r denotes Pearson correlation coefficient.

Next, we observed that *Mtb*-infected mice did not show a strong correlation (r=0.68) between pathogen load (as measured by colony forming units per organ) in the lungs and spleens (**Supplementary Fig. 2**) underscoring the heterogeneity of *Mtb* distribution in these organs of the humanized mice. This is consistent with an earlier report^41^ showing that increase of *Mtb* load in the lungs and spleens follow different trajectories over the course of infection. Therefore, we analyzed the correlation between metabolites abundances and *Mtb* load in spleens and lungs separately.

Interestingly, none of the metabolites correlated with the HIV load (shown in **Fig. 6c**) exhibited correlation either positively or negatively with *Mtb* load in lung or spleen. However, PC(16:0/18:1(11Z)) and lysoPC(16:0/0:0) positively correlated with *Mtb* load in lung as well as spleen (**Fig. 7b**). In addition, 3-hydroxyheptanoic acid exhibited a strong negative correlation with *Mtb* load in lung (**Fig. 7b**). Similarly, LysoPC(18:1/0:0) showed strong positive correlation, and myristic acid, PC(20:3(5Z,8Z,11Z)/20:3(5Z,8Z,11Z)) and PC(18:2(9Z,12Z)/16:0) showed strong negative correlation with *Mtb* load of the spleens (**Fig. 7c**).

## DISCUSSION

The development of animal models is a major requirement for developing drugs and vaccines for infectious diseases^42–44^. The lack of an ideal animal model can therefore delay the development of intervention strategies that can improve the outcome of disease in humans. The study of the interactions taking place during HIV/*Mtb* co-infection is particularly challenging due to a variety of factors, related to the nature of these pathogens, and the animal models. In this study, we demonstrated a reliable and reproducible small animal model for HIV/*Mtb* co-infection research using humanized NSG-SGM3 mice. We show that our model can recapitulate many aspects of HIV/*Mtb* co-infection in clinical settings, which will be helpful for characterizing the HIV*/Mtb*-induced immunopathogenesis, and to test therapeutics and vaccines.

A primary concern with using the mouse models for HIV/*Mtb* co-infection studies relates to the viral host range, which is naturally limited to humans and some NHPs^45, 46^. This poses restrictions on experimentation using NHPs, which require specialized infrastructure and personal training that is not widely available ^8^. However, this limitation has been circumvented to some extent by the use of immunocompromised mice strains that can engraft human stem cells and differentiate them into a variety of human immune cells, allowing for both HIV and *Mtb* infection and viral replication^14, 15, 18, 19, 47^. We show that the NSG-SGM3 mice allow stem cells to differentiate into a range of immune cells becoming susceptible to HIV infection and viral replication. This is due to the differentiation of human lymphoid lineage cell subsets, in particular generation of CD4^+^ T cells, which are the major target for HIV infection and replication. Moreover, the abundant differentiation of both lymphoid and myeloid lineage subsets allows for the assessment of immunological markers of disease relevance during HIV infection, and to measure vaccination-induced immune responses. A decreased CD4^+^/CD8^+^ ratio was observed in the humanized mice following HIV-1 infection, suggesting that our model reproduced similar immunological alterations observed during the natural infection of humans^48, 49^.

A comparative advantage that the NSG-SGM3 mice used in the present study over the previous generation of humanized mouse models is the transgenic expression of three human cytokine genes that enhance the differentiation and maturation of myeloid cell lineages and regulatory T cells^14^. This is particularly important, considering that these immune cells play important roles in controlling both HIV and *Mtb* growth and also serve as the target cells for these pathogens^50–53, 54, 55^. Moreover, the presence of granulomas, which are the hallmark of *Mtb* pathology, in the *Mtb*-infected humanized NSG-SGM3 mice is noteworthy, given that these structures are composed of multiple human immune cell populations from different lineages, that has not been seen in the C57BL/6 or BALB/c mice^56^. In addition, the previously reported humanized NSG-BLT mice required specialized surgical procedures in adult mice^19^, or the handling of newborns^14^. The humanization of NSG-SGM3 mice only requires a single intravenous injection of stem cells, which makes humanization much simpler to produce a viable small animal model for HIV/*Mtb* research.

We further note the differential expression of multiple human cytokines by the NSG-SGM3 humanized mice after HIV and *Mtb* single-infection or co-infection, which indicates that the reconstituted human immune cell subsets in these animals are functional and responsive during the infectious process. It should be noted that many of the cytokines that showed increased levels of expression in tissues after infection, were colony stimulating factors (G-CSF and GM-CSF) or chemoatractants (MCP-1, MIP-1α, MIP-1β), which have been implicated in human immune response against HIV and *Mtb*^57–62^. This indicates that immune cell recruitment and differentiation diverge according to the immune response induced by these pathogens in our model. Moreover, each tissue exhibited a different cytokine production profile. This could be due to the difference in cell types present in the tissues, as well as the viral/bacterial load and its effect on the immune response. In this regard, we noted that cytokine production did not increase in the lungs of the *Mtb* infection group, despite having a high bacterial load confirmed by culture. This is interesting and may suggest that *Mtb* suppresses lung immune responses to enhance its growth^52, 63–65^.

Nevertheless, cytokine expression in spleens was increased in the *Mtb*-infected mice, indicating immune activation in this organ. Similarly, the results of the Pearson correlation in plasma metabolites from the HIV-infected mice likely reflect the immune modulation by the pathogen, considering the positive correlation of viral load with an immunostimulatory xenobiotic (diethanolamine)^66^, while an inverse correlation was found with a subproduct of histamine metabolism (Imidazoleacetic acid)^67^. Although additional investigations are required, these results suggests concurrent activation of immune response, and suppression of the inflammation pathway; this coincides with earlier reports which show that histamine release is inversely correlated to the number of HIV-infected CD4+ T cells in humans^68^. The differences in cytokine and metabolite production may also reflect different stages of disease, and further studies are needed to validate these hypotheses.

The metabolome data also provided insight into the disruptions of the immunometabolism after HIV/*Mtb* infections in the humanized mice. It is noteworthy that the majority of the DAMs detected in the present study for both HIV and *Mtb* infection are fatty acids or metabolites involved in their metabolism. In accordance with previous reports, triglycerides were found to be increased in the plasma of HIV-infected mice, regardless of *Mtb* infection status^69^. Thus, Lysophosphatidylcholines (LysoPC), such as LysoPC (16:0/0:0), have been found to be increased in HIV-infected individuals^70^. Paradoxically, the concentration of palmitic acid (16:0), the fatty acid attached to the C-1 position of LysoPC (16:0/0:0), was found to be decreased in HIV-infected mice compared to the uninfected controls, suggesting a disruption in fatty acid metabolism. Moreover, dodecanoic (12:0), myristic (14:0) and arachidic (20:0) acids were also decreased in the HIV-infected mice, in line with a previous study that reported a reduction in free fatty acid concentration in serum from people living with HIV, which increased after antiretroviral treatment^71^. On the other hand, Pearson correlation showed an inverse relation between HIV load and imidazoleacetic acid, an imidazole receptor stimulator. Given the anti-HIV potential of the imidazole derivatives^72, 73^, the higher concentration of imidazoleacetic acid may facilitate the imidazole receptor binding, thus activating the imidazole-mediated anti-HIV capacity, and a lower HIV load. In addition, glucose metabolic pathways in regulating HIV infection in CD4+ T cells have been extensively reported^74, 75^. HIV infection increases glucose uptake in CD4+ T cells, and consequently, a higher glucose uptake by the CD4+ T cells will result in a lower concentration of glucose left in the serum; therefore, it was not surprising to see a negative correlation between HIV load and the metabolite glucose/mannose 6-phosphate in the serum (**Fig 7a)**.

In the case of *Mtb* infection, multiple DAMs related to TB pathogenesis were found in the plasma of infected mice (Table 2). Platelet-activating factor, increased in the *Mtb*-infected mice, has been previously shown to be an important part of TB immunopathology, and present in TB granulomas of humans and participating in the activation of other immune cell types during infection^76^. Meanwhile, N^6^-(Δ^2^-isopentenyl) adenine, a cytokinin previously thought to be produced only in plants, has been recently proven to be produced by *Mtb* (thus significantly increased in *Mtb*-infected mice), likely having a role in the protection of *Mtb* against nitric oxide^77^. Interestingly, three fatty acids (Dodecanoic acid, Myristic acid, and Palmitic acid) that were decreased in the HIV-infected mice were also decreased in plasma from *Mtb*-infected humanized mice, in addition to gluconic acid (6:0). The fatty acids alterations reflected the changes of mitochondrial function and β-oxidation, and this also is also evidenced by the reduction of L-carnitine, a metabolite necessary for the uptake of large chain fatty acids by the mitochondria^78^. We recall here that lipid-related metabolites have been reported to be decreased in humans co-infected with HIV and *Mtb*^79^. It has been reported that *Mtb* can alter lipid metabolism in macrophages, reducing the rate of ATP production, while at the same time, increasing their dependence on exogenous rather than endogenous fatty acids^80^. We therefore propose that the decrease of free fatty acids in the plasma of *Mtb*-infected animals might be related to sequestering of the pathogen in the macrophages^81^.

Collectively, our study shows that the NSG-SGM3 humanized mice can efficiently engraft human CD34+ stem cells which differentiate into a full lineage of functional immune cells. These mice are susceptible to both HIV-1 and *Mtb* infections, and the HIV/*Mtb* infections cause similar immunological, pathological, and metabolic changes in these mice as in humans. Therefore, the humanized NSG-SGM3 mice recapitulate the human-like immune responses to HIV/*Mtb* infections.

## Data availability

All data supporting the findings of this study are available in the manuscript. If there are any special requests or questions for the data, please contact the corresponding author (G.Y.).

## Acknowledgments

We thank Dr. Amy Tvinnereim for helping perform the following experiments: Irradiated the mice and performed *Mtb* infection of the humanized mice.

## Contributions

Guohua Yi: Conceived and guided the study, designed experiments, analyzed data, edited figures, and wrote and finalized the manuscript.

José Alejandro Bohórquez: Designed and performed the experiments, analyzed data, made the figures, and wrote the manuscript.

Sitaramaraju Adduri: Performed the experiments, analyzed data, made the figures, and wrote the manuscript.

Danish Ansari: Performed the experiments. Sahana John: Performed the experiments.

Jon Florence: Performed the experiments and analyzed data. Omoyeni Adejare: Performed the experiments.

Gaurav Singh: Performed the experiments. Nagarjun Konduru: Edited the manuscript.

Chinnaswamy Jagannath: Provided guidance on experimental design and edited the manuscript.

