## Supplementary materials for "A Novel Humanized Mouse Model for HIV and Tuberculosis Co-infection Studies"

**Supplementary Figure 1:** Gating strategy.

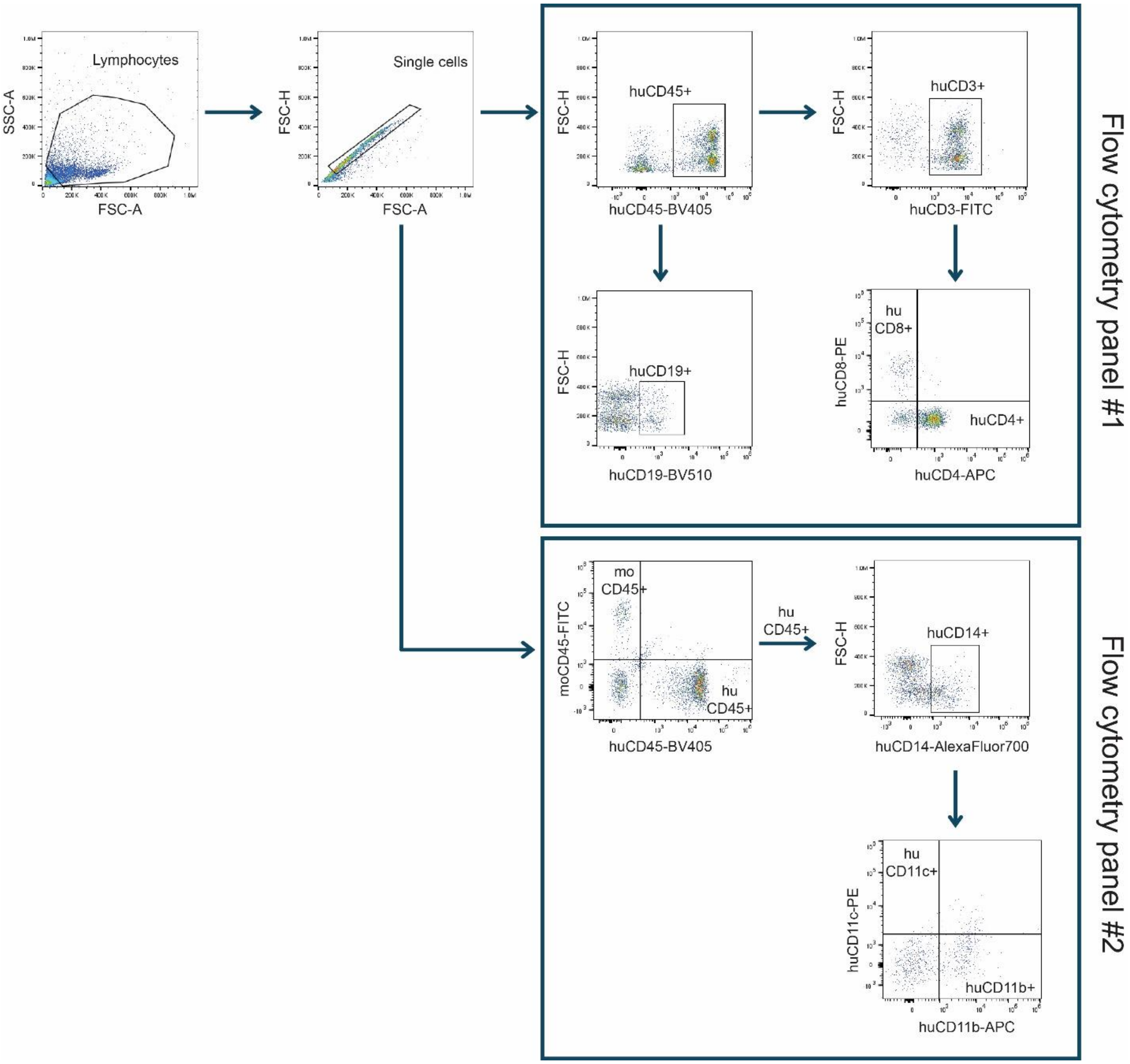

**Supplementary Figure 2:** Scatter plots showing Pearson correlation between Mtb load in mouse lungs and spleens (CFU/spleen).

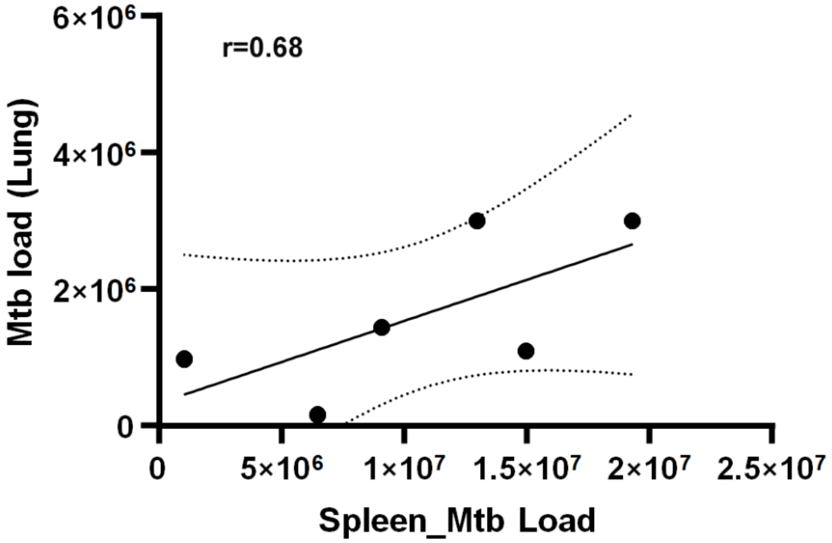

**Supplementary Table 1:** VIP scores computed for all the metabolites in the dataset based on OPLS-DA.

| **Metabolite** | **VIP score** |
|---|---|
| FA (16:0) | 1.437111 |
| N6 delta 2 isopentenyl adenine | 1.434205 |
| S-Adenosylmethionine | 1.418893 |
| FA (12:0) | 1.415167 |
| FA (18:0) | 1.396659 |
| Platelet-activating factor | 1.386854 |
| Lyso-PC (18:0) | 1.362849 |
| FA (14:0) | 1.350673 |
| FA (20:0) | 1.345078 |
| Deoxyuridine | 1.342263 |
| Carnitine | 1.33464 |
| Ascorbate | 1.308655 |
| 3-Hydroxybutyrate | 1.295623 |
| Tyrosine | 1.260681 |
| Indole-3-acrylate | 1.248672 |
| Lyso-PC (16:0) | 1.234492 |
| Indole-3-pyruvate | 1.218925 |
| 3-Hydroxycarnitine (5:0) | 1.218571 |
| Deoxycarnitine | 1.21358 |
| PE (38:3) | 1.208569 |
| Creatine | 1.206018 |
| Acetylcholine | 1.205549 |
| 2.Hydroxyglutarate.Citramalate | 1.201937 |
| N-alpha-Acetyllysine | 1.201507 |
| Gluconate | 1.197691 |
| Phenylalanine | 1.194949 |
| 4 Oxo-NAM | 1.187957 |
| Ribulose.Ribose.Xylulose_5.phosphate | 1.181456 |
| 6 Oxo-NAM | 1.179736 |
| FA (16:1) | 1.179513 |
| Pipecolate | 1.178893 |
| Dimethylglycine | 1.175609 |
| Pyridoxal | 1.174357 |
| Hexosamines | 1.170796 |
| Glucose | 1.170249 |
| PC (40:5) | 1.166408 |

| | |
|---|---|
| Tryptophan | 1.161525 |
| Glutarate.Monoethylmalonate | 1.15544 |
| Lyso-PC (17:0) | 1.153914 |
| Diethanolamine | 1.153088 |
| Alanine.beta.Alanine | 1.151022 |
| Adipate.Methyglutarate.Monomethylglutarate | 1.147097 |
| Suberate | 1.133155 |
| Isoleucine/Leucine | 1.131499 |
| .Fructose_6.phosphate.Galactose_1.phosphate | 1.127141 |
| Argininosuccinate | 1.125074 |
| Oxohexanoic acid | 1.124952 |
| Choline | 1.121099 |
| FA_8.0 Octanoic acid | 1.109269 |
| 3-Hydroxymethylglutarate | 1.10777 |
| FA (20:2) | 1.100887 |
| Glycerol_cyclic_phosphate | 1.099676 |
| Glycerophosphocholine | 1.096131 |
| Carnitinamide | 1.093773 |
| Asparagine | 1.092705 |
| Azelate | 1.092398 |
| Citrate.Isocitrate | 1.09206 |
| 3-(2-Hydroxyphenyl)propanoate | 1.088243 |
| Phosphoenolpyruvate | 1.077073 |
| Citrulline | 1.076909 |
| Aspartate | 1.071999 |
| Acetylacrylate | 1.070812 |
| Allothreonine.Homoserine.Threonine | 1.069962 |
| PC (40:6) | 1.066772 |
| PC (34:1) | 1.05803 |
| Lyso-PC (18:1) | 1.055564 |
| Hydroxy-FA (7:0) | 1.054677 |
| Lyso-PE (16:0) | 1.046704 |
| SM (d18:1/16:0) | 1.044607 |
| N-Acetylornithine | 1.035736 |
| PC (34:2) | 1.033627 |
| PE (38:4) | 1.031127 |
| Pentoses | 1.028832 |
| 3-Methoxy-4-hydroxymandelate | 1.024757 |
| 4-Imidazoleacetate | 1.022081 |
| N,N-Dimethylarginine | 1.020195 |
| Glucose_6.phosphate.Mannose_6.phosphate | 1.017318 |

| | |
|---|---|
| Ornithine | 1.017197 |
| PC (32:0) | 1.013559 |
| Pyruvate | 1.002992 |
| PC (38:4) | 0.999385 |
| Homoarginine | 0.999141 |
| Hexitols | 0.99771 |
| Methionine | 0.995206 |
| Proline | 0.989876 |
| 5.Aminolevulinate.cis..trans.3.Hydroxyproline.cis.4.Hydroxyproline.trans.4.Hydroxyproline | 0.981924 |
| PC (36:5) | 0.975882 |
| Glutamine | 0.974746 |
| 5,6 Dimethylbenzimidazole | 0.968548 |
| Lyso-PE (18:1) | 0.967866 |
| Carnitine (18:1) | 0.967755 |
| N,N-Dimethyllysine | 0.966673 |
| Carnitine (3:0) | 0.965 |
| Carnitine (16:1) | 0.963095 |
| PC (40:8) | 0.957938 |
| Galacturonate.Hexarates | 0.955387 |
| N,N,N-Trimethyllysine | 0.953082 |
| 3,4-Dihydroxyphenylacetate | 0.949528 |
| Urate | 0.945383 |
| Carnosine | 0.94116 |
| PC (38:5) | 0.940501 |
| 4alpha-Hydroxyformyl-4beta-methyl-5alpha-cholest-8-en-3beta-ol | 0.938381 |
| Stachyose | 0.938214 |
| PE (38:6) | 0.937361 |
| PC (36:2) | 0.935478 |
| trans-Aconitate | 0.935095 |
| FA (18:1) | 0.931431 |
| Aminocaprylate | 0.924364 |
| Tartrate | 0.921061 |
| N-Acetylaspartate | 0.920317 |
| Threonate | 0.914988 |
| PC (36:4) | 0.913867 |
| Lyso-PE (18:0) | 0.911883 |
| Serine | 0.90743 |
| 2',4'-Dihydroxyacetophenone/Resorcinol monoacetate | 0.906722 |
| Sulfolactate | 0.905885 |
| Glutathione | 0.905686 |
| Cystine | 0.901311 |

| | |
|---|---|
| PC (32:1) | 0.897269 |
| Arginine | 0.892206 |
| Diphthamide | 0.884566 |
| Raffinose | 0.882656 |
| Myoinositol | 0.877701 |
| Malate | 0.873186 |
| Urocanate | 0.868794 |
| Lyso-PC (18:2) | 0.867252 |
| Methyl 4-aminobutyrate | 0.865099 |
| 3-Hydroxycarnitine (4:0) | 0.863005 |
| Theanine | 0.855869 |
| Carnitine (16:0) | 0.849307 |
| Hydroxy-FA (10:0) | 0.849 |
| Quinate | 0.840404 |
| Sedoheptulose 7-phosphate | 0.83466 |
| Ureidopropionate | 0.829667 |
| SM (d18:1/18:0) | 0.828499 |
| Anserine | 0.824395 |
| Fumarate | 0.812664 |
| Uracil | 0.80356 |
| Oxoadipate | 0.79558 |
| Carnitine (glutaryl) | 0.788163 |
| 4-Guanidinobutanoate | 0.784761 |
| alpha-Ketoglutarate | 0.769371 |
| Methacholine | 0.769051 |
| Phosphorylcholine | 0.766295 |
| beta.Glycerophosphate.Glycerol_3.phosphate | 0.76434 |
| 2..3.Phosphoglycerate | 0.762515 |
| Hypotaurine | 0.762452 |
| PE (36:2) | 0.750665 |
| S-Adenosylhomocysteine | 0.746779 |
| Quinolinate | 0.743582 |
| Deoxyribose | 0.741554 |
| Carnitine (18:0) | 0.737904 |
| Lyso-PE (20:4) | 0.71974 |
| Guanidinosuccinate | 0.718122 |
| Taurine | 0.708903 |
| Fructose | 0.706283 |
| Lactate | 0.703849 |
| Carnitine (5:0) | 0.695629 |
| N-Acetylneuraminate | 0.693727 |

| | |
|---|---|
| Glutathione disulfide | 0.682004 |
| Cellobiose.Lactose.Maltose.Melibiose.Palatinose.Sucrose.Trehalose | 0.674089 |
| 2-Hydroxy-4-(methylthio)butanoate | 0.663318 |
| Glutamate | 0.656884 |
| Glu-Gln | 0.643373 |
| Hydroxy-FA (8:0) | 0.638001 |
| N-Acetylglutamate | 0.616066 |
| 2-Deoxyglucose | 0.610727 |
| Carnitine (4:0) | 0.591497 |
| PE (36:1) | 0.582848 |
| Stachydrine | 0.556746 |
| PC (32:2) | 0.547161 |
| Phosphoethanolamine | 0.541491 |
| Dihydroxyacetone phosphate | 0.455891 |
| Succinate | 0.436414 |
| N-Acetylserine | 0.274851 |
